# New Insights and Enhanced Human Norovirus Cultivation in Human Intestinal Enteroids

**DOI:** 10.1101/2020.11.12.380022

**Authors:** Khalil Ettayebi, Victoria R Tenge, Nicolas W. Cortes-Penfield, Sue E. Crawford, Frederick H. Neill, Xi-Lei Zeng, Xiaomin Yu, B. Vijayalakshmi Ayyar, Douglas Burrin, Sasirekha Ramani, Robert L. Atmar, Mary K. Estes

**Affiliations:** Department of Molecular Virology and Microbiology, Baylor College of Medicine (BCM), Houston, TX, USA; Department of Medicine, BCM, Houston, TX, USA; Section of Gastroenterology, Hematology and Nutrition, Department of Pediatrics, Baylor College of Medicine, and USDA/ARS Children’s Nutrition Research Center, Houston, TX, USA

## Abstract

Human noroviruses (HuNoVs) are the leading cause of epidemic and sporadic acute gastroenteritis worldwide. We previously demonstrated human intestinal stem cell-derived enteroids (HIEs) support cultivation of several HuNoV strains. However, HIEs did not support virus replication from every HuNoV-positive stool sample, which led us to test and optimize new media conditions, identify characteristics of stool samples that allow replication, and evaluate consistency of replication over time. Optimization of our HIE-HuNoV culture system has shown that: 1) A new HIE culture media made with conditioned medium from a single cell line and commercial media promote robust replication of HuNoV strains that replicated poorly in HIEs grown in our original culture media made with conditioned media from 3 separate cell lines; 2) GI.1, eleven GII genotypes (GII.1, GII.2, GII.3, GII.4, GII.6, GII.7, GII.8, GII.12, GII.13, GII.14 and GII.17) and six GII.4 variants, can be cultivated in HIEs; 3) successful replication is more likely with virus in stools with higher virus titers; 4) GII.4_Sydney_2012 virus replication was reproducible over three years; and 5) HuNoV infection is restricted to the small intestine, based on replication in duodenal and ileal HIEs but not colonoids from the same donors. These results improve the HIE culture system for HuNoV replication. Use of HIEs by several laboratories worldwide to study the molecular mechanisms that regulate HuNoV replication confirms the usefulness of this culture system and our optimized methods for virus replication will advance the development of effective therapies and methods for virus control.

**Importance:** Human noroviruses (HuNoVs) are highly contagious and cause acute and sporadic diarrheal illness in all age groups. In addition, chronic infections occur in immunocompromised cancer and transplant patients. These viruses are antigenically and genetically diverse and there are strain-specific differences in binding to cellular attachment factors. In addition, new discoveries are being made on strain-specific differences in virus entry and replication and the epithelial cell response to infection in human intestinal enteroids. Human intestinal enteroids are a biologically-relevant model to study HuNoVs; however, not all strains can be cultivated at this time. A complete understanding of HuNoV biology thus requires cultivation conditions that will allow the replication of multiple strains. We report optimization of HuNoV cultivation in human intestinal enteroid cultures to increase the numbers of cultivatable strains and the magnitude of replication, which is critical for testing antivirals, neutralizing antibodies and methods of virus inactivation.

## Introduction

Noroviruses, members of a genus in the *Caliciviridae* family, are nonenveloped positive-sense RNA viruses of approximately 7.4-7.7 kb, and are classified based on sequence similarities into ten genogroups, amongst which genogroups GI, GII, GIV, VIII and IX infect humans (1). Within these five genogroups, there are 39 different genotypes that cause human infections; GIs and GIIs that are the most prevalent are divided into 9 and 27 genotypes (including eight GII.4 variants), respectively (1). Human noroviruses (HuNoVs) are a leading cause of diarrheal illness and are associated with nearly 20% of all gastroenteritis episodes worldwide, including 19-21 million cases and 56,000-71,000 hospitalizations annually in the United States (2, 3). The spectrum of illness ranges from acute morbidity in all age groups, chronic disease in immunocompromised cancer and transplant patients, and mortality in young children and older adults (2, 4–7). It is estimated that 685 million episodes of acute gastroenteritis and 212,000 deaths occur worldwide due to HuNoVs each year (8). Since the introduction of rotavirus vaccines, HuNoVs have become the leading cause of acute gastroenteritis in children worldwide (2, 9, 10). The economic burden of HuNoV gastroenteritis is substantial, with over $4 billion in healthcare costs and over $60 billion in societal costs annually (5). These data emphasize the strong need for effective therapies, antivirals and vaccines.

The lack of a reproducible culture system for HuNoVs was a major barrier to understanding virus biology including mechanisms of replication, inactivation, neutralization, and vaccine development for approximately five decades (11, 12). This problem was overcome with the successful cultivation of multiple HuNoV strains in enterocytes in human intestinal stem cell-derived, nontransformed enteroid (HIE) monolayer cultures (12–20). Previous studies showed replication of GI.1 and six GII genotypes, including four GII.4 variants, in this ex vivo system, and virus replication in HIEs mimics epidemiological differences in host susceptibility based on genetic differences in expression of histo-blood group antigens (HBGAs) as defined by a person’s secretor status (11–13, 21, 22). In addition to being used to study the regulation of viral replication and pathophysiology, the HIE cultivation system allows the evaluation of antiviral candidates, neutralization and methods for virus inactivation (12, 13, 15, 17, 20, 23, 24). Despite this progress, a need to improve the system remained, primarily because not every HuNoV-positive stool sample could be propagated in HIEs.

Significant advancement has been made in the development and maintenance of ex vivo long-term HIE cultures since they were originally established in 2011 (25). Growth factors, including R-spondin, Wnt-3A, and Noggin, are needed to support vital pathways for stem cell maintenance (26). Due to the high cost and reduced biologic activity of some purified commercial growth factors, these factors often are made by expression individually or in combination in mammalian cell lines, where they undergo posttranslational modifications prior to their secretion into the culture media to produce conditioned medium. This conditioned medium is filtered and used as growth factor supplements in HIE proliferation or expansion medium to sustain the maintenance of multicellular three dimensional (3D) cultures, which then are used to produce monolayer (2D) cultures for infection experiments where access to the apical surface is needed (27). Components in the conditioned media, such as serum or other factors produced by the cultures expressing the growth factors, may positively or negatively affect HIE growth and/or viral infection. In this study, we sought to assess the reproducibility of HuNoV infections in HIEs over time, optimize the *ex-vivo* HIE system for HuNoV replication in order to increase the numbers of cultivatable strains and the magnitude of replication, and identify factors that result in successful virus replication.

## Results

### Many HuNoV strains replicate in jejunal J2 HIE monolayers

We previously reported the establishment of the HIE system for HuNoV cultivation and demonstrated the replication of GI.1, GII.3, GII.17 and 4 GII.4 variants (12). Here, we tested additional stools representing a greater spectrum of HuNoV strains to evaluate whether they can be propagated in the ex vivo HIE culture system and to examine culture conditions that affect virus growth. J2 HIE monolayers cultivated and plated in our original in-house proliferation (BCMp) media and then differentiated in our in-house differentiation (BCMd) media were inoculated with HuNoV-positive fecal filtrates, and virus replication was assessed by RT-qPCR using GI.1 or GII.4 transcripts for quantification of genome equivalents (GEs). A 0.5 log_10_ increase in GEs after 24 hpi relative to the amount of genomic RNA detected at 1 hpi (after removal of the virus inoculum and two washes of the monolayers to remove unbound virus) was set as a threshold to indicate successful viral replication. In total, virus replication was seen in 31/40 stool samples tested, including one GI genotype (GI.1) and eleven GII genotypes (GII.1, GII.2, GII.3, GII.4, GII.6, GII.7, GII.8, GII.12, GII.13, GII.14, GII.17). The GII.4 samples included six variants (GII.4_2002, Yerseke_2006a, Den Haag_2006b, New Orleans_2009, Sydney_2012, and Sydney_2015) (Table 1). Increases in HuNoV GEs at 24 hpi ranged from 0.5-3.38 log_10_. Viruses in stool samples that did not grow in the J2 HIEs included GI.3, and 8 GII.4 strains.

**Table 1:**
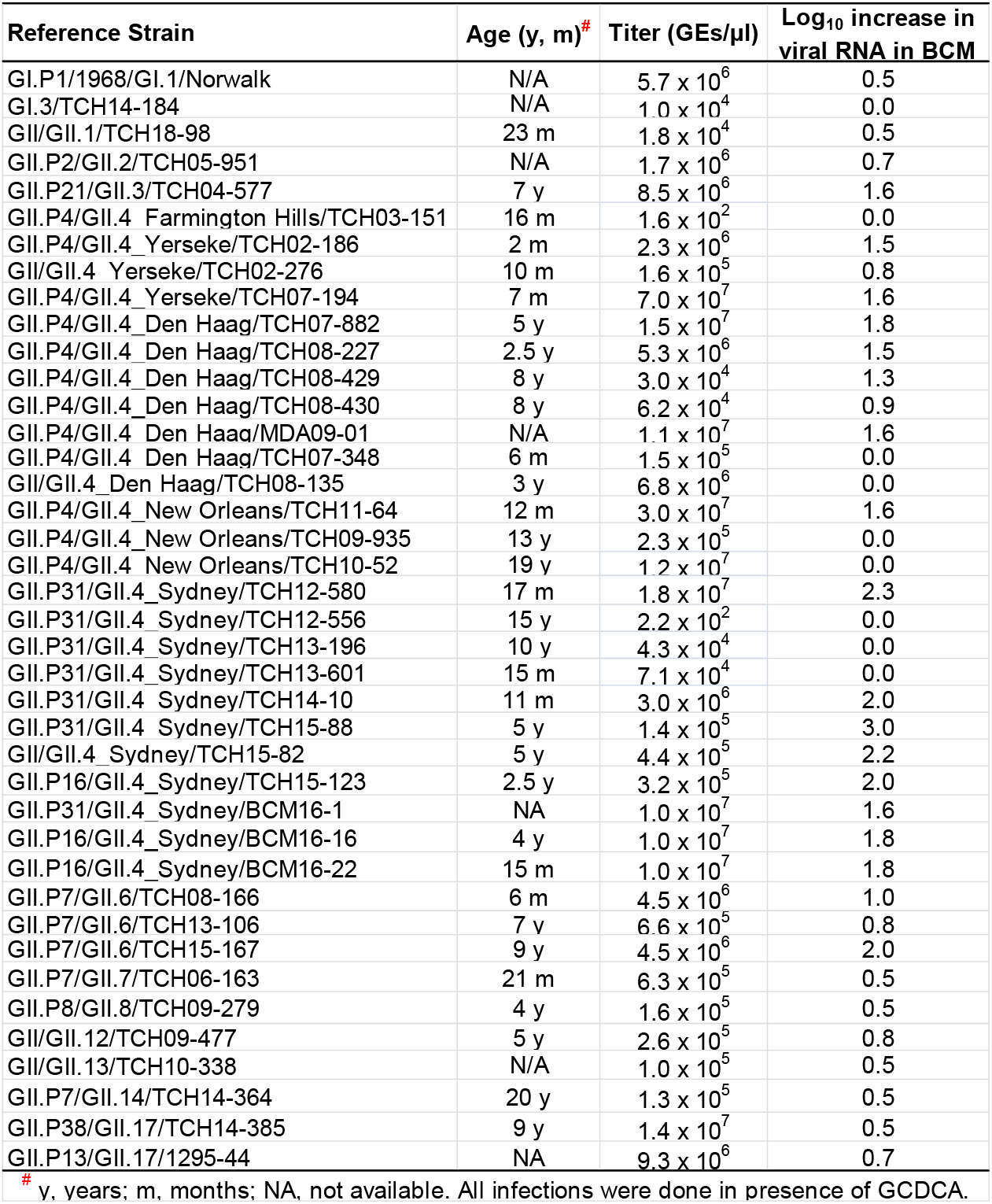
HuNoV strains tested for cultivation in jejunal HIEs cultured in BCM media

### Successful replication is more likely with virus from stools with higher virus titers

To examine factors that may affect HuNoV infection, we assessed differences in replication based on stool viral load and age of infected person. Previously, we determined the infectious dose required for replication of GII.4_Sydney_2012 in 50% of cultures (TCID_50_) to be ~1200 GEs/well. We found that replication of GII.4 strains was more likely to occur with fecal samples from patients with a viral titer greater than 1200 GEs per μL (Fig 1A). Replication was not affected by patient age (Fig. 1B).

**FIG 1.**
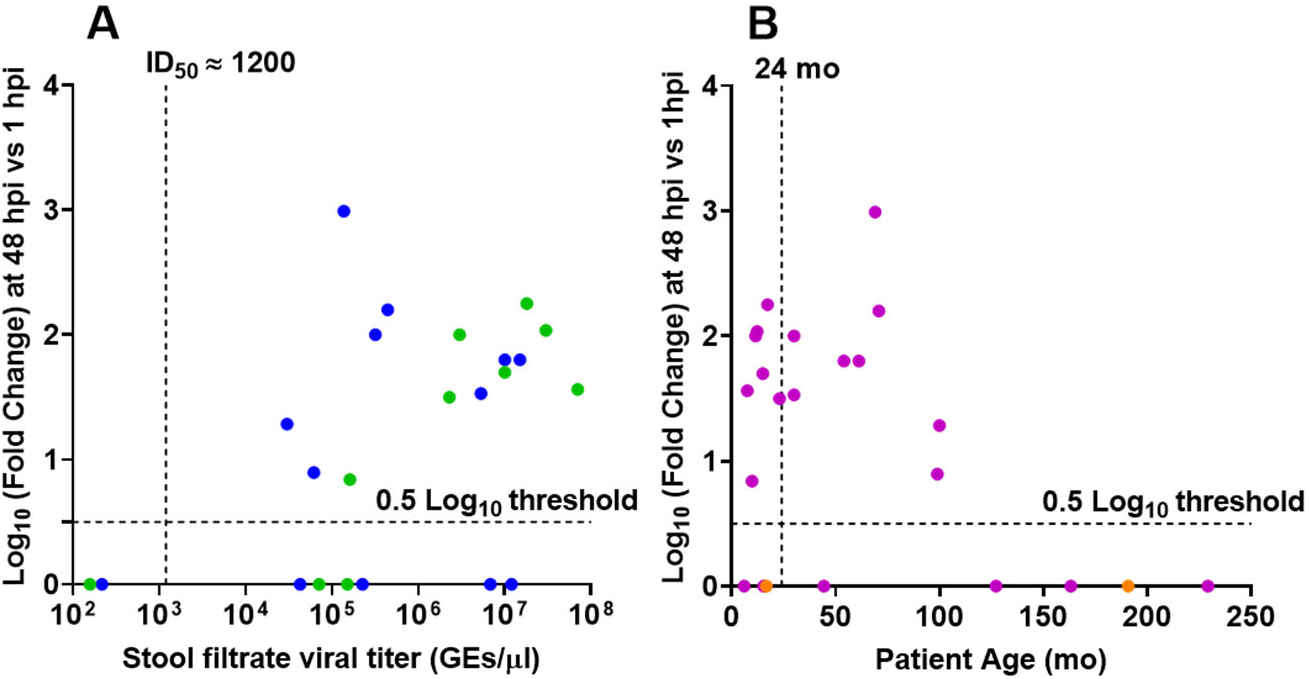
Successful replication is more likely with virus from stools with higher virus titers (A) but not affected by patient age (B). (A) Replication of GII.4 strains plotted by virus titer. Dashed vertical line indicates the GII.4_Sydney_2012 ID_50_ determined previously (12). Data points indicate stools from patients under (green) and over (blue) 2 years of age. (B) Replication of the same set of GII.4 strains plotted by patient age. Purple, stool titer > GII.4_Sydney_2012 ID_50_. Orange, stool titer < GII.4_Sydney_2012 ID_50_. Replication less than 0.5 log_10_ was assigned a value of 0. Dashed lines show the detection 0.5 log_10_ threshold.

We previously reported from our human challenge studies with GI.1 virus that higher prechallenge levels of HuNoV-specific fecal IgA correlated with a reduced peak viral load (28). We set out to determine whether fecal antibody levels correlated with the ability to cultivate GII.4 HuNoVs. Filtrates were prepared from 12 stools from pediatric patients with GII.4 norovirus gastroenteritis (age range 7.5 mo – 19 yr), and total and GII.4-specific IgA levels from each stool filtrate were quantified. Although total IgA was detected in each of the stool filtrates, GII.4-specific IgA was not detected in any of the samples (Table S1).

**Table S1:** IgA levels in GII.4 HuNoV-positive stool filtrates

| Strain | Age of Patient | IgA present? | Total IgA (ug/ml) | GII.4-specific IgA present? | Infectivity ($\text{Log}_{10}$ increase in viral replication in BCM medium at 24 hpi vs 1 hpi ) |
| --- | --- | --- | --- | --- | --- |
| TCH07-194 | 7 m | Y | 3.4 | N | 1.6 |
| TCH07-882 | 5 y | Y | 11.8 | N | 1.8 |
| TCH08-135 | 3 y | Y | 5.6 | N | 0 |
| TCH08-227 | 2 y | Y | 2.9 | N | 1.5 |
| TCH09-935 | 13 y | Y | 0.9 | N | 0 |
| TCH10-52 | 19 y | Y | 1.4 | N | 0 |
| TCH11-64 | 12 m | Y | 7.8 | N | 1.6 |
| TCH12-556 | 15 y | Y | 11.1 | N | 0 |
| TCH13-196 | 10 y | Y | 11.5 | N | 0 |
| TCH14-10 | 11 m | Y | 76.4 | N | 2.0 |
| TCH15-123 | 2 y | Y | 9.2 | N | 2.0 |
| TCH15-82 | 5 y | Y | 12.4 | N | 2.2 |

Based on these results, it was not possible to draw conclusions whether infectivity is associated with levels of HuNoV-specific IgA in fecal samples, but the lack of replication in the five non-cultivable samples could not be attributed to the presence of such IgA.

### HuNoV replication is reproducible over time

We also assessed the reproducibility of the HIE system by comparing the efficiency of GII.4 replication in our standard HIE cultivation media used in previous HuNoV studies (proliferation in BCMp and differentiation in BCMd media) over a three year period (2016-2019). A GII.4_Sydney_2012 virus was included as a positive control in all experiments conducted in our laboratory over the 3-year period, and although there was variability in viral yields over time, virus replication occurred consistently (Fig. 2). The geometric mean virus increase at 24 hpi was 2.26 log_10_ (n=80; SD=0.4), with 3.38 log_10_ as the highest observed change.

**FIG 2.**
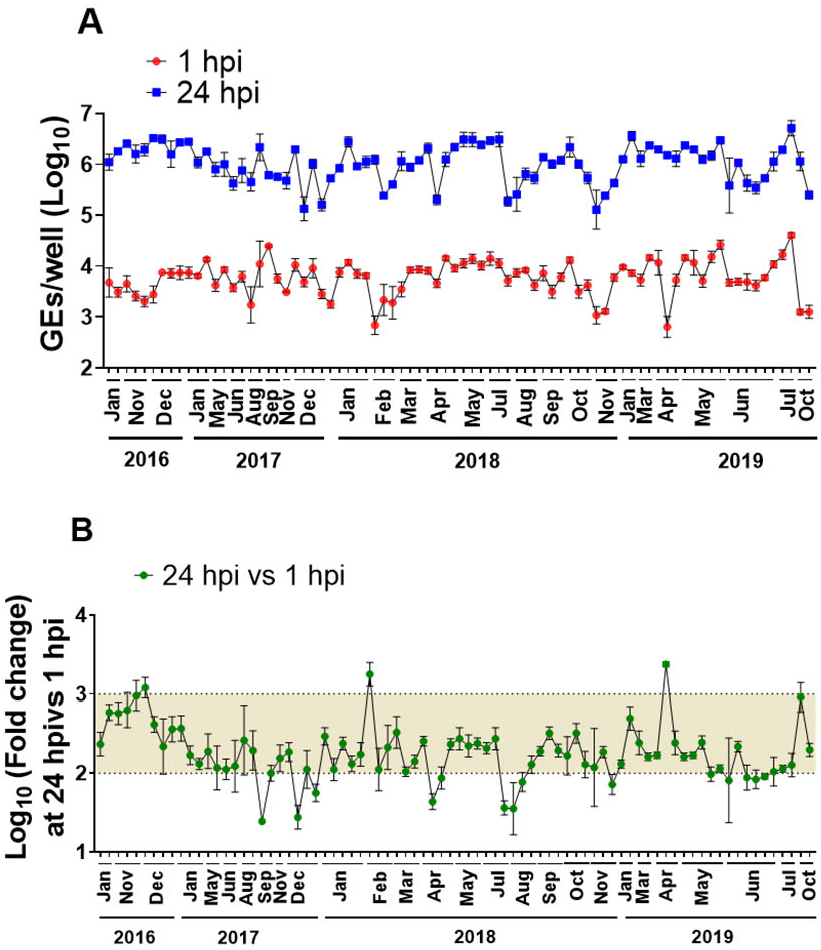
Replication of GII.4_2012_Sydney in HIEs plated in BCM medium is reproducible over time. (A) Virus replication of GII.4_Sydney_2012 HuNoV, included as a positive control in different experiments throughout 3+ years to assess the reproducibility of viral infection in J2 HIE monolayers infected with 9 × 10^5^ GEs/well in BCM media, was determined at 1 hpi and 24 hpi. (B) Fold change at 24 hpi compared to 1 hpi. The mean log_10_ increase at 24 hpi versus 1 hpi was 2.25 (n=80). Error bars denote standard deviation from 6 wells in each experiment.

### HuNoVs replicate efficiently in HIEs cultured in Intesticult medium

To further evaluate and potentially simplify cultivation conditions, we next compared GII.4_Sydney_2012 HuNoV replication in jejunal HIE monolayers plated in BCMp medium and differentiated with BCMd medium to replication in HIEs plated in a commercially available medium [Intesticult (referred here to as INTp and INTd) human organoid growth medium from Stem Cell Technologies]. At 24 hpi, the geometric mean log_10_ GE increases (Δ24hpi-1hpi) were significantly higher in four different jejunal HIE monolayers plated in INT media compared to BCM media (Fig. 3A). Similar results were obtained with another HuNoV genotype (GII.3, Fig. 3B), suggesting that using the commercially available INT media to plate jejunal HIE lines efficiently promotes better replication of HuNoV strains compared to BCM media.

**FIG 3.**
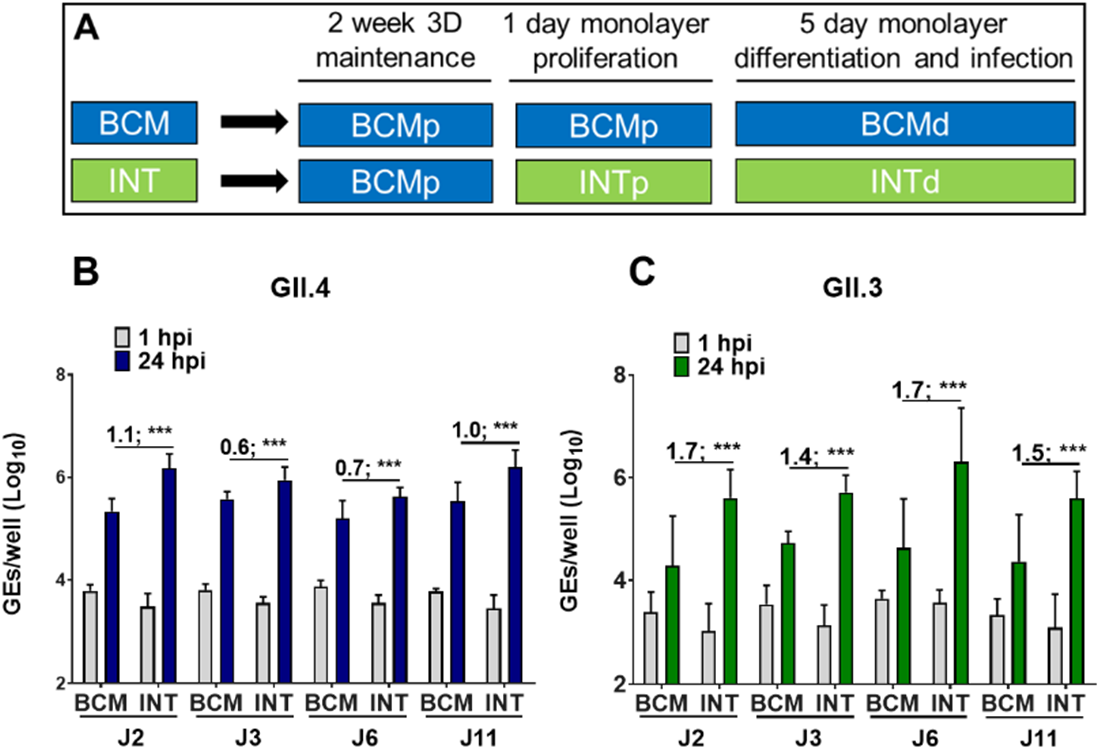
Improved HuNoV replication in different jejunal HIE cultures plated as monolayers in INT media. (A) Schematic design of HIE culture maintenance and monolayer seeding prior to infection (“p” and “d” notations refer to proliferation and differentiation respectively). (B) GII.4_Sydney_2012 (9 × 10^5^ GEs/well) and (C) GII.3 (4.3 × 10^5^ GEs/well) virus replication was evaluated in four jejunal (J2, J3, J6 and J11) HIE lines plated in either BCM or INT media. Each experiment was performed twice and compiled data are presented. Error bars denote standard deviation (N=12). Values on the bars indicate log_10_ (fold change) replication difference in BCM vs BCM media at 24 hpi. Significance was determined using Student’s t-test(***, p value < 0.001).

We previously reported that HIEs derived from the three segments (duodenum, jejunum, and ileum) of the small intestine support HuNoV replication when cultured in BCM media and that enterocytes are the primary target for infection and replication (12, 14). However, since these lines were derived from different donors, it was not possible to directly examine segment-specific differences in susceptibility in the absence of confounding genetic differences between individuals. We have now established HIE cultures from three intestinal segments (duodenum, ileum, and colon) from two secretor-positive donors (104 and 109) (29). Duodenal, ileal and colonic HIEs from donors 104 and 109 were plated in BCMp or INTp media, and after differentiation, were inoculated with GII.4_Sydney_2012 or GII.3 HuNoVs in the corresponding differentiation media (Fig. 4).

**FIG 4.**
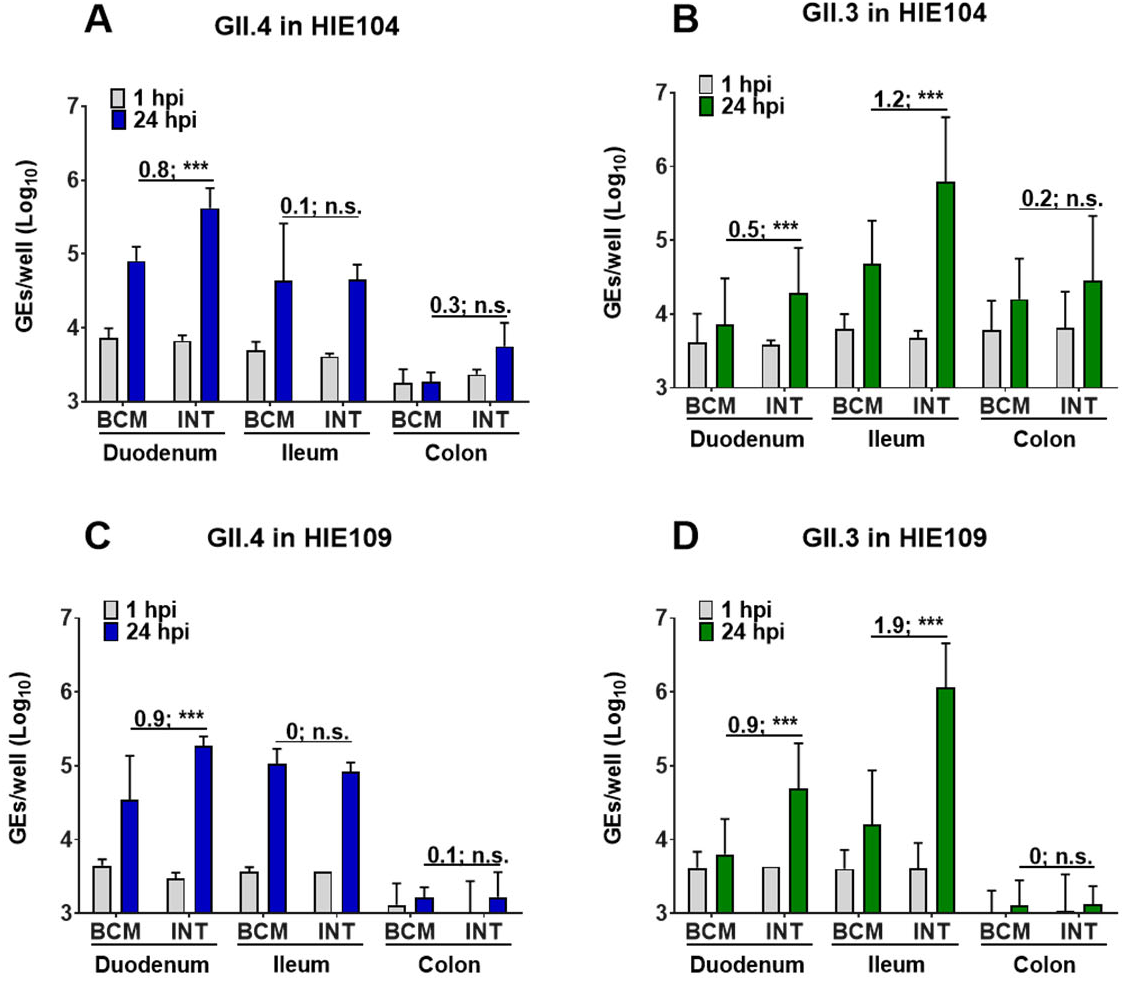
HuNoV replication in HIE cultures from different intestinal segments (duodenum, ileum, colon) from two independent donors (104 and 109). HIEs were plated in BCMp or INTp media (see schematic design in FIG 3A). After differentiation, monolayers were infected with (A, C) GII.4_Sydney_2012 (9 × 10^5^ GEs/well) or (B, D) GII.3 (4.3 × 10^5^ GEs/well). Compiled data from two experiments are presented. Error bars denote standard deviation (n=12) and each data bar represents the mean for six wells of inoculated HIE monolayers. Values on the bars indicate log_10_ (fold change) replication difference in INT vs BCM media at 24 hpi. Significance was determined using Student’s t-test (***, p value < 0.001; n.s., not significant).

Replication of GII.4_Sydney_2012 was observed in duodenal HIE monolayers from both donors, with significantly greater GE increases (Δ24hpi-1hpi) when monolayers were plated and differentiated in INT media compared to BCM media (mean of 0.8 and 0.9 log_10_ increases in HIE 104 and HIE 109, respectively) (Fig. 4A, 4C). Duodenal HIEs did not support GII.3 replication when cultured in BCM media, but geometric mean increases of 0.7 log_10_ and 0.9 log_10_, respectively, were attained in INT media for donors 104 and 109 (Fig. 4B, 4D). Both GII.4_Sydney_2012 and GII.3 viruses replicated in ileal HIE monolayers; however, while INT media promoted greater GII.3 replication compared to BCM media, there was no difference in replication for GII.4_Sydney_2012. Colonic HIEs did not support GII.4_Sydney_2012 and GII.3 replication when cultured in either medium.

We previously showed bile acids induce multiple cellular responses that promote GII.3 replication, and J2 HIEs do not support GII.3 replication when cultured in BCMd medium without addition of the bile acid glycochenodeoxycholic acid (GCDCA) (24). Since replication of GII.3 is consistently higher in INTd medium (Fig. 3 and 4), we investigated whether INTd medium promotes GII.3 replication in the absence of GCDCA and whether the addition of GCDCA further enhances GII.3 replication. As expected, no significant GII.3 replication was detected in BCMd medium without GCDCA. In INTd medium, a 0.9 log_10_ increase in GEs (Δ24hpi-1hpi) was seen in the absence of GCDCA and 1.3 log_10_ increase in GEs was seen at 24 hpi in the presence of GCDCA (Fig. S1). In the presence of GCDCA, a 0.4 log_10_ greater increase in GEs (Δ24hpi-1hpi) was seen in INTd compared to BCMd media. These results suggest that INT media may contain component(s) that promote GII.3 virus infection and act synergistically in the presence of GCDCA to enhance GII.3 replication.

We next investigated whether INT media enhanced viral replication of other HuNoV strains. Almost all (20/21) HuNoV positive stool samples representing both GI and GII viruses, replicated as well or significantly better (18/21) in HIEs plated in INT media compared to BCM media (Table 2).

**FIG S1.**
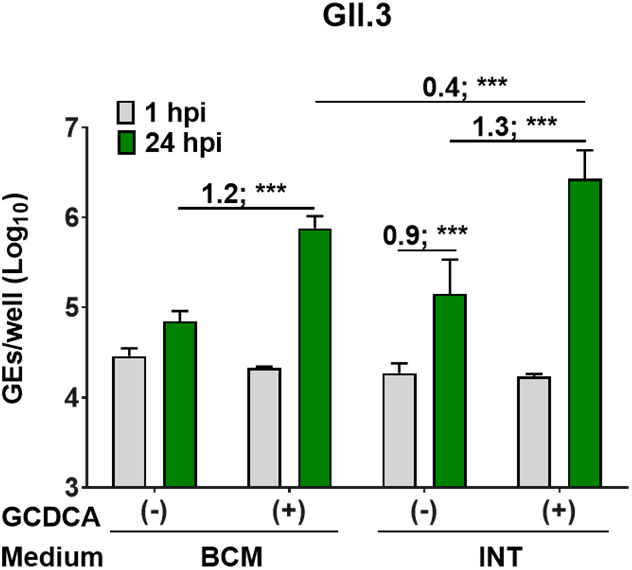
HIE cultures plated in INT media support some GII.3 replication in J2 HIE monolayers and replication is further enhanced by addition of GCDCA supplement. 3D J2 HIEs were maintained in BCM proliferation medium. Monolayers were proliferated and differentiated in INT or BCM media (see schematic design in FIG 3A), and inoculated with GII.3 (4.3 × 10^5^ GEs/well) diluted in CMGF(−) with or without 500 μM GCDCA supplement. After 1 hpi, monolayers were washed twice and cultured in the indicated differentiation medium. Values above bars represent Log_10_ difference in viral growth at 24 hpi vs 1 hpi. Values represent the mean and error bars denote standard deviation (n=6). Asterisks indicate significant difference from INT medium at 24 hpi: *** p value < 0.001.

**Table 2:** HuNoV strains successfully replicated in jejunal HIEs plated in BCM vs INT media

| Reference Strain | Titer<br>(GEs/μl) | Log<sub>10</sub> increase<br>in viral RNA in<br>BCM medium | Log<sub>10</sub> increase in<br>viral RNA in INT<br>medium |
| --- | --- | --- | --- |
| GI.P1/1968/GI.1/Norwalk | 5.7 x 10<sup>6</sup> | 0.5 | 1.9 * |
| GI.3/TCH14-184 | 1.0 x 10<sup>4</sup> | 0.0 | 0.0 |
| GII/GII.1/TCH18-98 | 1.8 x 10<sup>4</sup> | 0.5 | 1.7 * |
| GII.P2/GII.2/TCH05-951 | 1.7x 10<sup>6</sup> | 0.7 | 2.1 * |
| GII.P21/GII.3/TCH04-577 | 8.5 x 10<sup>6</sup> | 1.6 | 2.1 * |
| GII.P4/GII.4_Yerseke/TCH07-194 | 7.0 x 10<sup>7</sup> | 1.5 | 2.0 * |
| GII.P4/GII.4_Den Haag/TCH07-882 | 1.5 x 10<sup>7</sup> | 1.8 | 2.2 * |
| GII.P4/GII.4_Den Haag/MDA09-01 | 1.1 x 10<sup>7</sup> | 1.6 | 2.4 * |
| GII.P4/GII.4_New Orleans/TCH11-64 | 3.0 x 10<sup>7</sup> | 1.6 | 1.7 |
| GII.P31/GII.4_Sydney/TCH12-580 | 1.8 x 10<sup>7</sup> | 2.3 | 2.7 * |
| GII.P16/GII.4_Sydney/BCM16-16 | 1.0 x 10<sup>7</sup> | 1.8 | 2.2 * |
| GII/GII.4_Sydney/TCH15-82 | 4.4 x 10<sup>5</sup> | 2.2 | 2.8 * |
| GII.P31/GII.4_Sydney/BCM16-1 | 1.0 x 10<sup>7</sup> | 1.6 | 2.0 * |
| GII.P7/GII.6/TCH13-106 | 6.6 x 10<sup>5</sup> | 0.8 | 1.3 * |
| GII.P7/GII.6/TCH15-167 | 4.5 x 10<sup>6</sup> | 2.0 | 2.9 * |
| GII.P8/GII.8/TCH09-279 | 1.6 x 10<sup>5</sup> | 0.5 | 0.7 * |
| GII/GII.12/TCH09-477 | 2.6 x 10<sup>5</sup> | 0.8 | 1.1 |
| GII/GII.13/TCH10-338 | 1.0 x 10<sup>5</sup> | 0.5 | 0.9 * |
| GII.P7/GII.14/TCH14-364 | 1.3 x 10<sup>5</sup> | 0.5 | 1.5 * |
| GII.P38/GII.17/TCH14-385 | 1.4 x 10<sup>7</sup> | 0.5 | 2.1 * |
| GII.P13/GII.17/1295-44 | 9.3 x 10<sup>6</sup> | 0.7 | 1.5 * |

\*, significant increase in INT vs BCM media. All infections were done in presence of GCDCA

We next examined the reproducibility of viral replication in J2 HIE monolayers plated in INTp medium and differentiated in INTd medium supplemented with GCDCA over a 12-month period (Fig. 5). GII.4_Sydney_2012 showed significant increases in viral GEs at 24 hpi vs 1 hpi with a mean of 2.66 log_10_ (n = 24), which is a 0.4 log_10_ increase compared to the reproducibility of GII.4 replication in BCMd medium throughout 3 years (Fig. 2).

**FIG 5.**
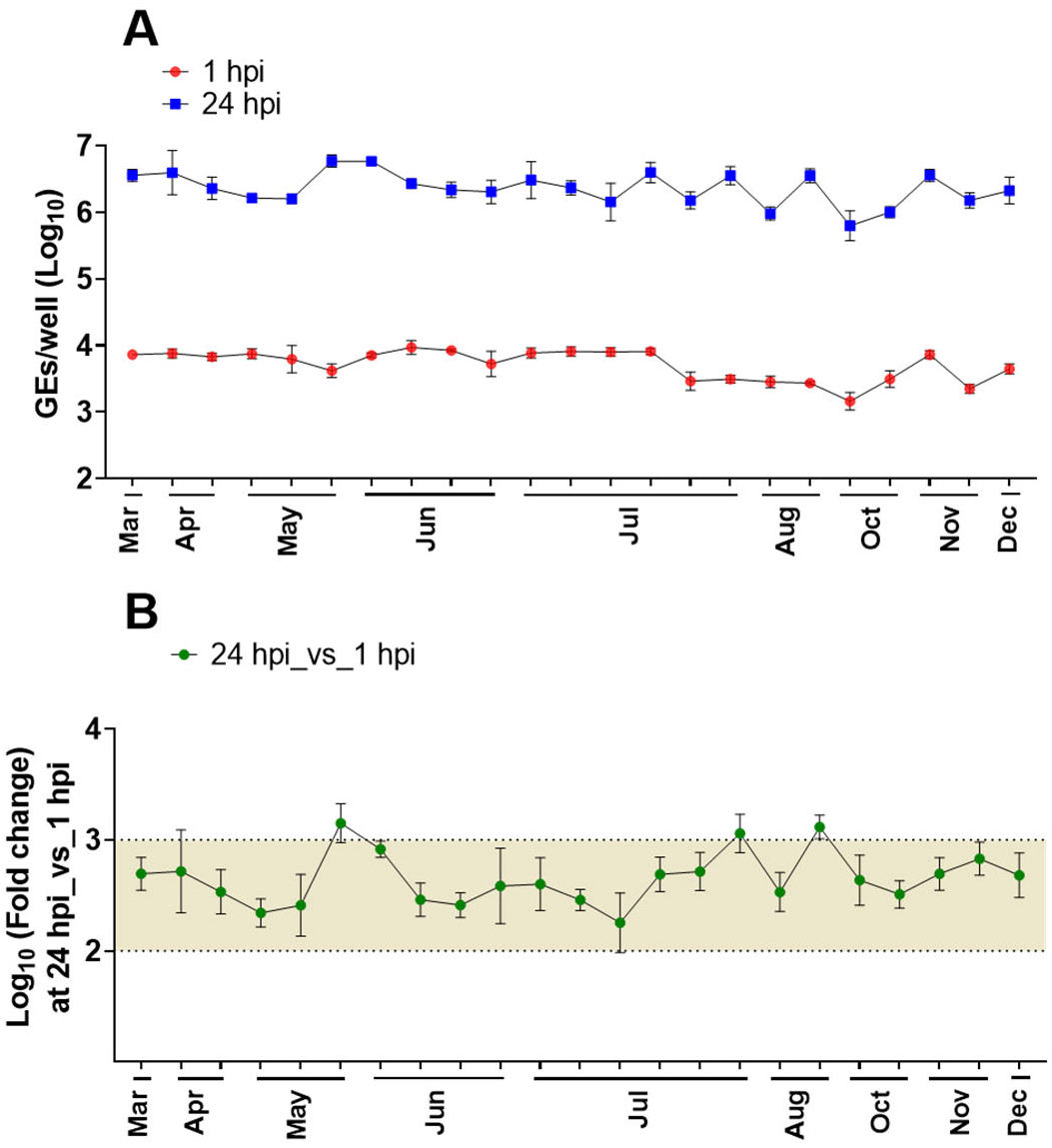
Replication of GII.4_2012_Sydney in HIEs plated in Intesticult medium is reproducible and less variable over one year. (A) Virus replication of GII.4_2012_Sydney HuNoV, included as a positive control in different experiments throughout 1 year (2019) to assess the reproducibility of viral infection in J2 HIE monolayers inoculated with 9 × 10^5^ GEs/well] in INT media was determined at 1 hpi and 24 hpi. (B) Fold change at 24 hpi compared to 1 hpi. The mean log_10_ increase at 24 hpi versus 1 hpi was 2.66 (n=23). Error bars denote standard deviation (n=6).

While we were completing studies comparing HuNoV replication in BCM and INT media, an ATCC L-WRN cell line (ATCC CRL3276) engineered to co-express and secrete the three growth factors (Wnt-3A, R-Spondin, and Noggin) became available (30, 31). This propagation media, referred to as L-WRN (described in material and methods section), uses a single cell line to produce the growth factors to make proliferation medium and offers several advantages in terms of reducing production time and effort compared to our original method of making the growth factors in three separate cell lines to make BCMp medium. Therefore, we assessed HuNoV replication in J2 HIEs propagated in BCMp or L-WRN media and then plated and differentiated in BCM or INT media. Replication of GII viruses from 4 genotypes was enhanced in J2 HIEs propagated in BCMp medium and then plated and differentiated in INT media (BCM/INT media) compared to replication in HIEs propagated, plated and differentiated in BCM media (BCM/BCM media) (Fig. 6), consistent with our previous results (Fig. 3). Similar results were obtained when J2 HIEs were propagated in L-WRN medium and then plated and differentiated in either BCM or INT media (L-WRN/INT vs L-WRN/BCM) (Fig. 6). Moreover, enhancement of replication of GII.4_Sydney_2012 and GII.17 was significantly greater in the L-WRN/INT combination compared to BCM/INT (Fig. 6B and 6D). In contrast, no significant differences in virus replication were observed for GII.3 and GII.6 replication in L-WRN/INT compared to BCM/INT.

**FIG 6.**
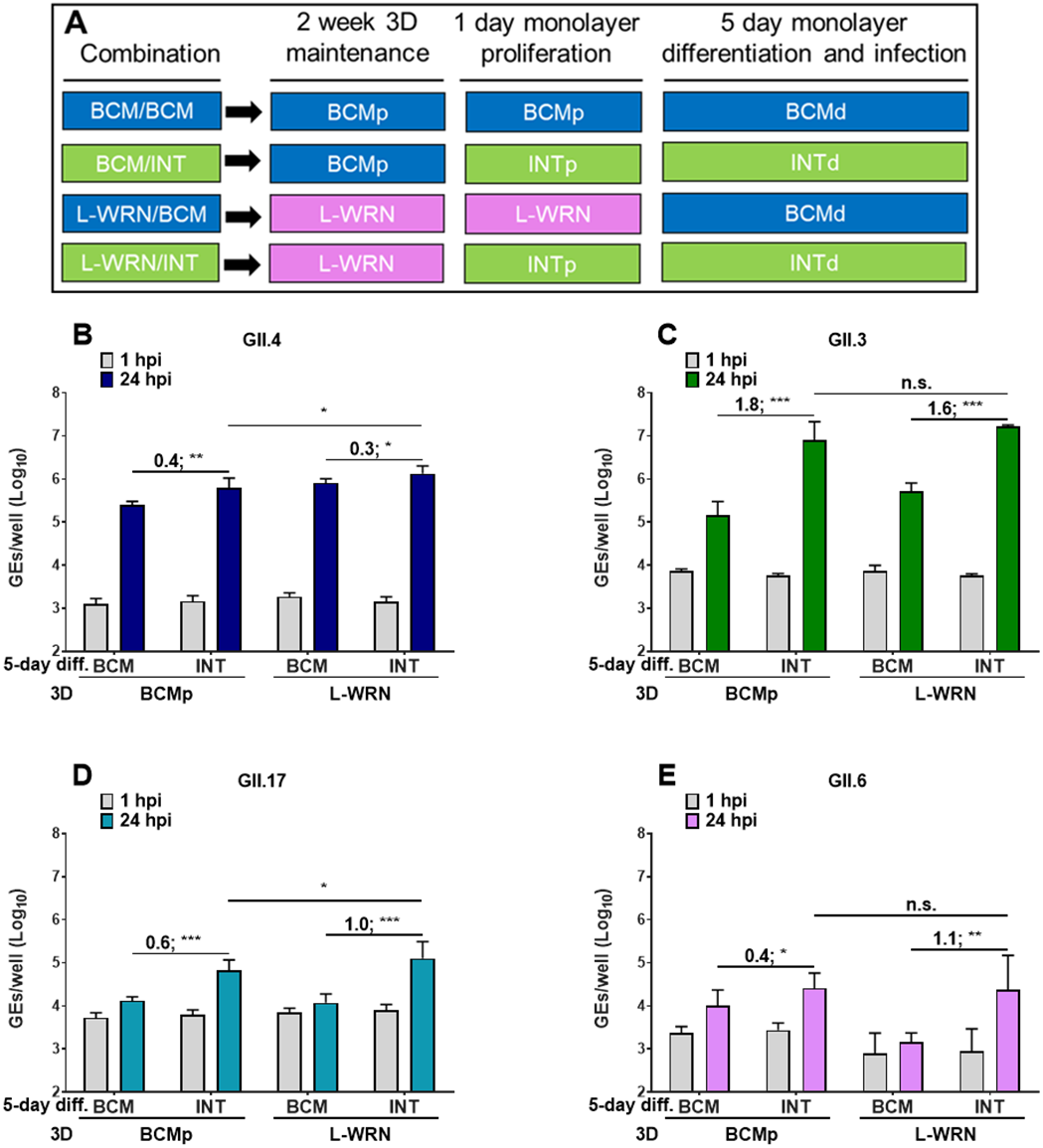
Replication of different GII HuNoV genotypes is improved when HIEs are plated in INT media. J2 HIEs were initially maintained as 3D-HIEs in BCMp or L-WRN proliferation media and then plated and differentiated for 5 days in the indicated media. (A, experimental design). Monolayers were inoculated with (B) GII.4_Sydney_2012 (9 × 10^5^ GEs/well), (C) GII.3 (4.3 × 10^5^ GEs/well), (D) GII.17_1295-44 (4.6 × 10^5^ GEs/well) or (E) GII.6_TCH13-106 (3.3 × 10^5^ GEs/well) diluted in CMGF(−) with 500 μM GCDCA. After 1 hpi, monolayers were washed and cultured in the indicated BCM or INT differentiation medium (+ 500 μM GCDCA). Values represent Log_10_ difference in viral growth between conditions [(Δ24hr-1 hr in INT) – (Δ24hr-1 hr in BCM)]. Error bars denote standard deviation (n=6). Asterisks indicate significant difference between conditions. *** p value < 0.001; ** p < 0.01; * p < 0.05; n.s., not significant.

To further investigate the difference between INT and L-WRN media with regard to HuNoV replication, we compared the replication of GII.4_Sydney_2012, GII.3, GII.17, and GII.6 in monolayers prepared from 3D J2 HIEs propagated for two weeks in either INTp or L-WRN media. Monolayers were plated from INT-3D- or L-WRN-3D-HIEs, proliferated for one day and then differentiated for 5 days in the indicated media (Fig. 7). The highest levels of replication of GII.4_Sydney_2012, GII.3 and GII.17 were observed when INT media was used for J2 HIE propagation, plating and differentiation (Fig. 7A, 7B and 7C). When propagating 3D J2 HIEs in L-WRN medium and plating/differentiating monolayers in INT media (L-WRN/INT combination), viral replication was significantly increased (0.6, 1.0, 0.3, 0.6 log_10_ increases for GII.4_Sydney_2012, GII.3, GII.17 and GII.6, respectively) compared to the L-WRN/BCM media combination. Virus replication remained significantly reduced for GII.4_Sydney_2012 and GII.3 (0.5 and 0.4 log_10_ decreases, respectively) in the L-WRN/INT combination compared to the INT/INT combination. Thus, while the INT medium for propagating, plating and differentiating HIEs was the best combination to support HuNoV replication, propagating HIEs in L-WRN and seeding monolayers in INT medium also achieved efficient HuNoV infections; this L-WRN/INT combination is attractive because it is more cost effective than using INT medium for both propagating and plating cells.

**FIG 7.**
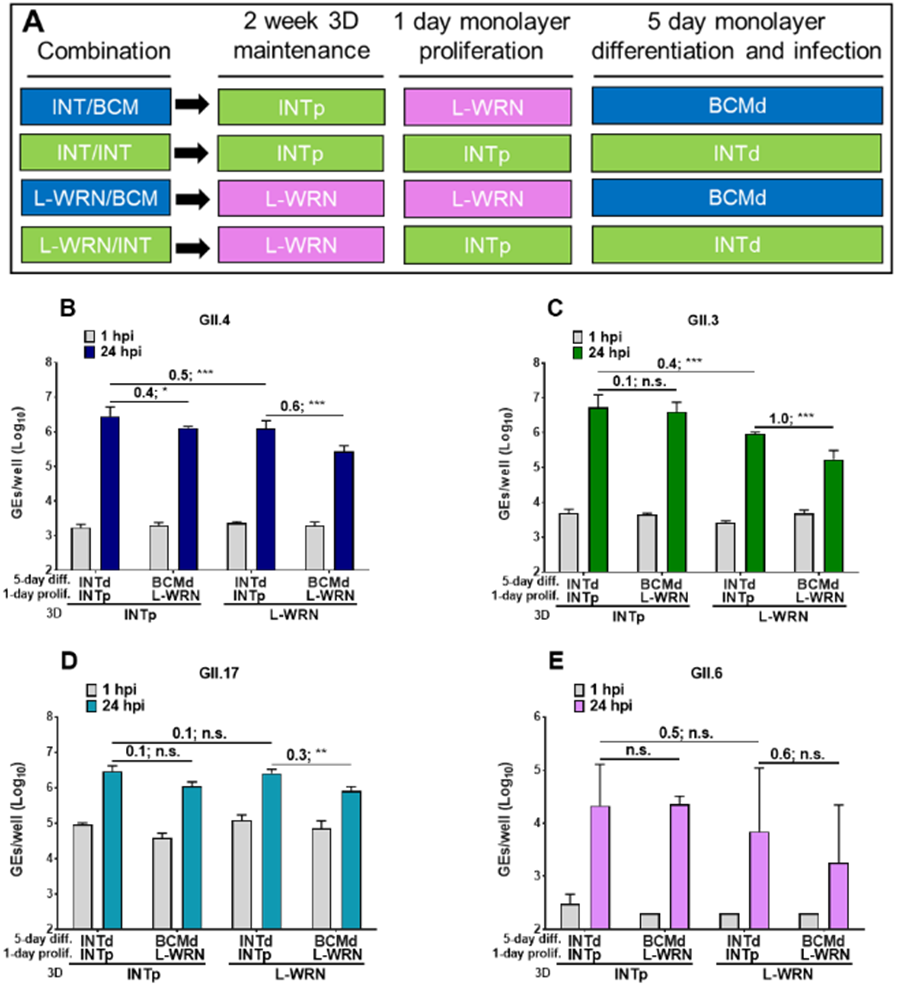
Media composition can improve HuNoV replication. J2 HIEs were propagated in INT or L-WRN media prior to seeding them into monolayers. Monolayers were prepared from INT-3D- or L-WRN-3D-HIEs, proliferated for one day and then differentiated for 5 days in the indicated media (A, experimental design). They were inoculated with (B) GII.4_Sydney_2012 (9 × 10^5^ GEs/well), (C) GII.3 (4.3 × 10^5^ GEs/well), (D) GII.17_1295-44 (4.6 × 10^5^ GEs/well) or (E) GII.6_TCH13-106 (3.3 × 10^5^ GEs/well) diluted in CMGF(−) with 500 μM GCDCA. After 1 hpi, monolayers were washed twice and cultured in the indicated differentiation medium (+ 500 μM GCDCA). Values above bars represent Log_10_ difference in viral growth between conditions. Error bars denote standard deviation (n=6). Asterisks indicate significant difference between conditions.

HuNoVs infect differentiated enterocytes (12, 14). To determine whether cultivation in the different media tested in this study affects differentiation, we next evaluated J2 HIEs for the gene expression of cell proliferation/differentiation markers after the cultures were grown in each proliferation medium (BCMp, INTp, L-WRN) and then differentiated in the corresponding differentiation medium (BCMd or INTd). Differentiation markers for enterocytes (sucrase isomaltase, *SI)*; goblet cells (mucin, *MUC2)*; and Paneth cells (alkaline phosphatase, *AP)* were significantly expressed in HIEs maintained in all media conditions, while gene expression for the stem cells markers (*LGR5*, *CD44*) and proliferation marker (*KI67*) were reduced (Fig. S2). Thus, HIE monolayers cultured for 5 days in either differentiation medium were fully differentiated, with no significant differences in enterocyte marker SI between the three conditions.

**FIG S2.**
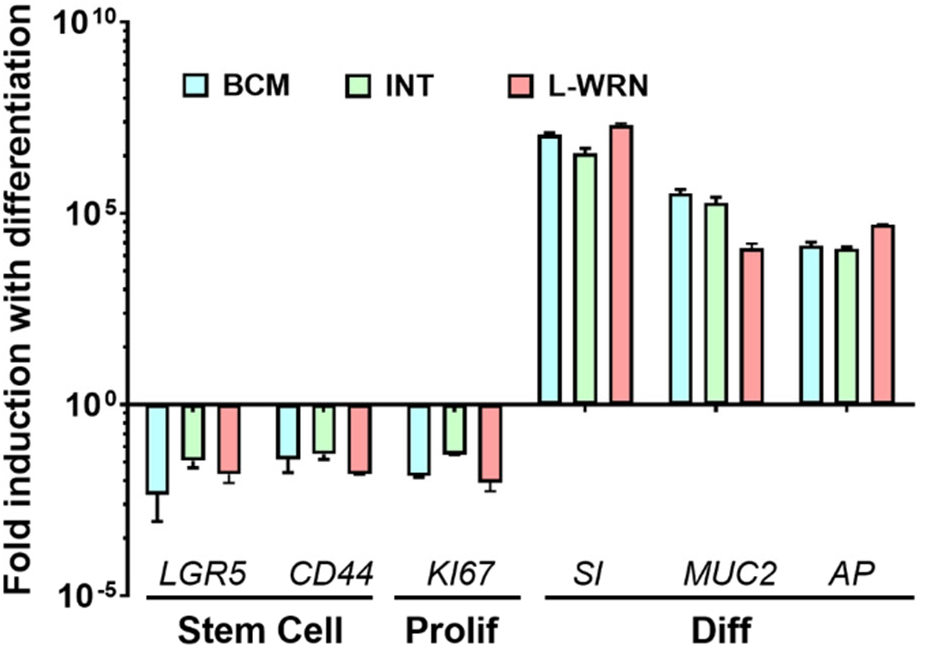
Media composition does not change differentiation of HIEs. Fold change in levels of transcripts assessed by RT-qPCR in differentiated J2 HIE monolayers relative to the transcript levels in proliferating J2 HIE monolayers. Transcript levels were first normalized to GAPDH levels prior to obtaining the relative fold change by using the 2^−ΔΔCT^ method. Shown are markers for stem cells, proliferating and differentiated cells. Gene symbols represent leucine-rich-repeat-containing G-protein-coupled receptor 5 *(LGR5),* antigen identified by monoclonal antibody Ki-67 *(K/67)*, CD44, sucrase-isomaltase *(SI)*, alkaline phosphatase (*AP*), and mucin 2 *(MUC2)* genes. Error bars indicate standard errors of the means (n = 3).

We are interested in further exploring how the L-WRN and INT media improve HuNoV replication. This is difficult to determine because the composition of INT media is proprietary. Based on our previous studies demonstrating increased replication of HuNoV strains in the presence of bile acids, we hypothesized that serum in INT media contains bile acids, which enhances virus replication (12, 21, 24). We analyzed INT components A (used in differentiation media) and B (used in combination with component A for proliferation media) as well as BCMp and L-WRN media for the presence of 12 different bile acids. We focused our analysis primarily on GCDCA and TCDCA (taurochenodeoxycholic acid) that facilitates GII.3 infection of HIEs at 5 μM and higher concentrations (24). Bile acids were not detected in INT component A. While INT component B had detectable levels of individual bile acids, none surpassed 5 μM (Fig. S3A). The concentration of bile acids in L-WRN media (sum=1.26 uM) was more similar to INT component B (sum=1.65 uM) than to INT component A or BCMp media. Since the source of bile acid in in-house media (BCMp and L-WRN) is likely to be the fetal bovine serum (FBS) used in the cultivation of growth factor cell lines, we next tested the bile acid concentrations in the different FBS lots used in our laboratory. For BCMp, Corning FBS was used at 8% while 10% Hyclone FBS was used in the L-WRN media. While both FBS contain GCDCA and TCDCA, the sum concentrations are almost double in the FBS used for L-WRN media compared to the one used for BCM media (Fig. S3B). The differences in the source and concentrations of FBS used, and corresponding differences in bile acid concentration may all provide explanations for the enhancement of HuNoV replication in HIEs grown in L-WRN and INT media compared to BCM media. Whether other factors in each of these media that affect HIE metabolism or health remains unknown.

**FIG S3.**
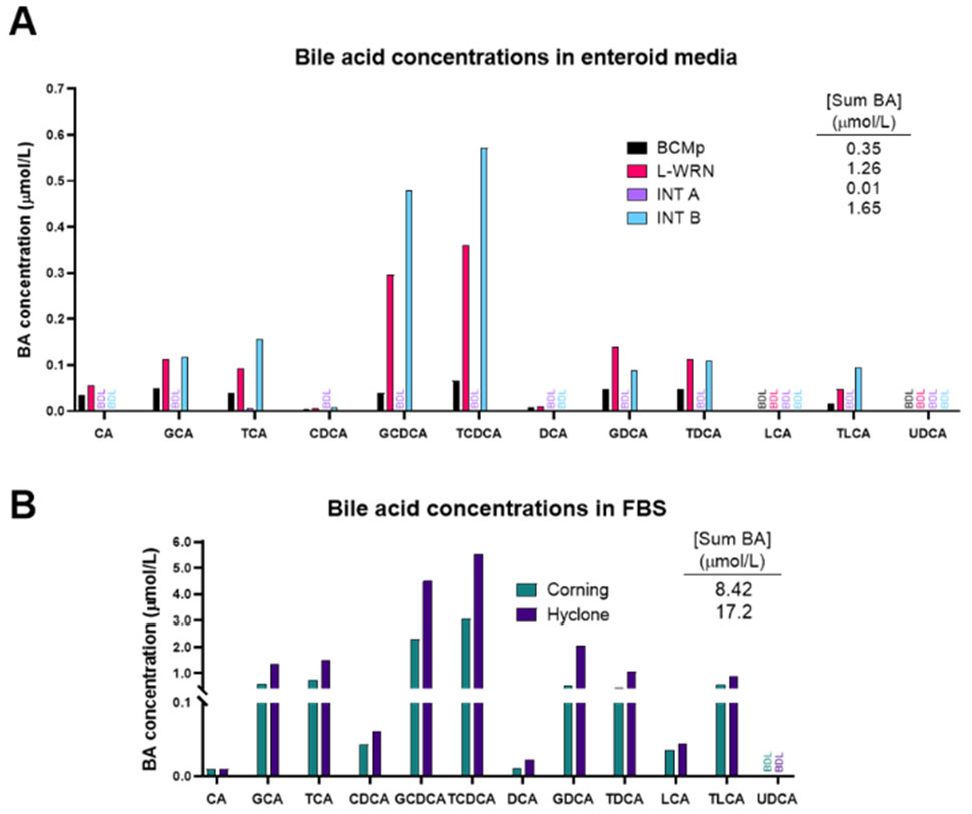
Enteroid media and FBS contain bile acids at low levels. (A) media [BCMp, L-WRN, INT component A (INT A) and INT component B (INT B)] or (B) commercial FBS (Corning or Hyclone brand) were analyzed by mass spectrometry (MS) for concentrations of a panel of individual bile acids. Inset tables contain the additive concentration of tested bile acids detected by MS (Sum BA).

## Discussion

After the HIE system was established for HuNoV cultivation, major advances have ensued in the norovirus field, manifested by studies that have focused on transcriptomic analysis, detecting viral infectivity, developing virus inactivation and neutralization methods, and dissecting the strain-specific requirement for bile for GII.3 entry for viral replication (13, 15–17, 24, 32, 33). In this manuscript, we report studies that centered on improving the HIE system to enhance viral replication and expand the spectrum of cultivatable HuNoV strains. We and others previously demonstrated successful replication of GI.1 and six GII genotypes (GII.1, GII.2, GII.3, four GII.4 variants, GII.14, and GII.17) in J2 HIEs (12, 13). Here we expanded the spectrum of cultivatable strains to cover five more genotypes (GII.6, GII.7, GII.8, GII.12 and GII.13) and two more GII.4 variants (GII.4_2002 and GII.4_Sydney_2015) (Tables 1 and 2). We found varied increases in virus yields when infections were performed in HIE monolayers cultured in BCM media. However, INT media enhanced replication of several strains belonging to different genotypes. Indeed, significant increases in virus yield were achieved for 18 strains tested in INT media compared to BCM media (Table 2). To our knowledge, this is the first study comparing the commercial media to laboratory-produced media related to HuNoV growth. Moreover, while reproducible replication of GII.4_Sydney_2012 strain was seen in either BCM or INT media, assessment of replication in INT media over a one year period showed higher fold increases in GEs and lesser variability, with a 2.66 log_10_ (95% CI 2.56-2.76; n= 23) mean increase in GEs at 24 hpi vs 1 hpi in INT media compared to 2.26 log_10_ (95% CI 2.17-2.33; n= 80) in BCM media (Figs. 2 and 5). While improvement in replication using the INT media is a major advance, the cost of this commercial medium is an issue for large scale culturing of HIEs. Indeed, we cannot afford to grow all our cultures in INT medium. Our final results using L-WRN medium for culturing HIEs followed by plating the cells in INTp medium prior to differentiation and infection in INTd medium, are important in identifying conditions that are cost-effective and still achieve enhanced replication of multiple viral strains. We and others have also previously described other methods of achieving enhanced HuNoV replication such as through genetic modification of HIE fucosylation (GI.1 and GII.17) or of HIE innate responses (GII.3) (16, 21). Culturing genetically modified HIEs in these optimized culture media conditions may further optimize the HIE-HuNoV culture system and expand the spectrum of cultivatable GI genotypes.

Replication of GI.1, GII.1, GII.3, GII.6 and GII.17 in BCM media are bile acid-dependent (data not shown), providing another example of strain-specific differences in requirements for HuNoV replication (12, 18, 21, 24). We previously defined the BA-mediated mechanism for GII.3 replication involves virus uptake mediated by dynamic and rapid BA-mediated cellular endolysosomal dynamic changes and cellular ceramide (24). Future studies are necessary to confirm whether the uptake and subsequent replication of all these BA-dependent strains are regulated by the same mechanism as shown for GII.3.

Costantini et al. reported successful HuNoV replication in HIEs was largely contingent on initial virus titer and genotype, even though several GII genotype samples with moderate or high viral titers failed to replicate (13). While HuNoV infection is primarily restricted based on secretor status of the cultures, GII.3 has unusual characteristics (12, 21, 24). This raises the question of whether there are other unknown factors for successful replication of noncultivatable strains. Previously, Costantini et al. (13) tested 80 stools containing various HuNoV genotypes from young patients (0-12 years old) or adults over 18 and found that 15/16 replicating strains came from stools from the 0-12 year age group. They further showed that 13/16 replicating strains were from patients under two years old. Of the 25 GII.4 strains we tested, all replicating strains were isolated from stools of patients under 12. However, we found no statistical difference in replication of strains derived from patients under 2 years of age compared to stools from older patients. High viral titer is one predictor of successful replication and inoculating cultures with a dose that is above the minimal infectious dose is the best predictor of replication success (12, 13, 17). However, six among eight GII.4 strains that did not show positive replication had moderate to high titers (4.3 × 10^4^ – 1.1 × 10^7^ GEs/μL) (Fig. 1A and Table 1). The reasons related to failure to achieve viral replication of strains with high titers remain unclear but could be due to the presence of noninfectious particles in the fecal sample, cellular host factors or anti-HuNoV antibodies in the fecal samples that might impede the binding of viral particles to their specific surface receptors. We previously showed that pre-challenge levels of Norwalk-specific fecal IgA correlates with reduced viral load in human experimental infection studies (34, 35). However, we were unable to detect norovirus-specific IgA in stools where virus did not grow.

While many advances have been made to generate 3D human organoids, different media compositions are used in the context of the specificity and function of the original tissue. The media formulations and growth factors are considered vital elements required for efficient establishment and long-term maintenance of organoids. Due to the high cost of commercial purified growth factors, most organoid media, including that for HIEs, are formulated with growth factors expressed in eukaryotic cell lines and supplemented as conditioned media. These conditioned media retain impurities, such as serum components and possibly bile acids, that may positively or negatively affect HIE growth and/or viral infection. We showed that HIE media and FBS present in conditioned media contain bile acids at low levels, which may explain the enhancement of HuNoV replication in HIEs grown in L-WRN and INT media.

Here, we evaluated the effect of ex vivo HIE growth on HuNoV infectivity in two laboratory-produced versus commercially available media: BCM (designated in previous studies as CMGF+), the first medium published by Clevers et al. (25) and previously used in HuNoV replication system (12–14); L-WRN formulated with one conditioned media that has three growth factors expressed from one cell line (36); and the commercial INT. With regards to HuNoV infectivity, INT and L-WRN media both promoted GII.3, GII.4_Sydney_2012, GII.6 and GII.17 replication better than BCM media when HIEs are maintained and differentiated in INT media (Fig. 7). Due to the high cost of the commercial media, we have found a simplified cost-effective way to use a combination of L-WRN/INT, by sustaining the 3D HIE growth in L-WRN and seeding and infecting HIE monolayers in INT media.

## Material and methods

### Preparation of HuNoV positive/negative stool filtrates

Ten percent stool filtrates containing HuNoV were prepared as described previously (12). In brief, 4.5 mL of ice-cold PBS was added to 0.5 mL of stool, homogenized by vortexing, and sonicated three times for 1 min. The sonicated suspension was centrifuged at 1,500 × *g* for 10 min at 4°C. The supernatant was transferred to a new tube and centrifuged a second time. The resulting supernatant was passed serially through 5 μm, 1.2 μm, 0.8 μm, 0.45 μm and 0.22 μm filters depending on stool texture. The filtered sample was aliquoted and frozen at −80°C until used (Table 1).

### Human intestinal enteroid culture

All HIE cultures used in this study are from an HIE bank maintained by the Texas Medical Center Digestive Diseases Center (TMC DDC) Core. Jejunal HIE cultures were previously generated from surgical specimens obtained during bariatric surgery. Duodenal, ileal, and colonic HIE cultures were generated from biopsy specimens obtained from adults during routine endoscopy at Baylor College of Medicine (BCM) through the TMC DDC Study Design and Clinical Research Core. The BCM Institutional Review Board approved the study protocols. All HIEs used in this study were secretor-positive (Table S2).

HIEs were obtained from the DDC Core as multilobular cultures in Matrigel. HIEs were maintained and propagated in 24-well plates as previously described (12, 14). Monolayer cultures in 96-well plates were prepared for infection from the multilobular cultures. Each well of a 96-well plate was coated with 33 μg/mL collagen IV diluted in 100 μl ice-cold water that was removed between a minimum of 2 hours and a maximum of overnight incubation at 37°C. Multilobular HIEs were washed with 0.5 mM EDTA in ice cold DPBS (calcium chloride-magnesium chloride free) and dissociated with 0.05% trypsin/0.5 mM EDTA for 5 min at 37°C. Trypsin was then inactivated by adding CMGF[-] medium (12) supplemented with 10% FBS to the cell suspension. Cells were dissociated by pipetting 50 times with a P1000 pipet and passing them through a 40 μm cell strainer. Cells were pelleted for 5 min at 400 × *g*, suspended in proliferation medium containing the ROCK inhibitor Y-27632 (10 μM, Sigma), and seeded onto a 96-well plate at a concentration of 100,000 cells per well. After 1 day of cell growth as a monolayer, the proliferation medium was changed to differentiation medium. The cells were maintained in the differentiation medium for 5 days, with the medium being changed every other day.

### Media

Five different media were used to maintain and differentiate HIEs:

1. Complete medium with growth factors (BCMp), prepared at BCM by the DDC core, consisted of CMGF[−] medium supplemented with epidermal growth factor (EGF), nicotinamide, gastrin I, A-83-01, SB202190, B27 supplement, N2 supplement, *N*-acetylcysteine, and Noggin, R-spondin, and Wnt3A conditioned media prepared from three different expressing cell lines (12, 37). Noggin- and R-spondin-expressing cell lines were kindly provided by Van den Brink (Amsterdam, The Netherlands) and Calvin Kuo (Palo, CA, USA), respectively. L-Wnt-3A-expressing cell line (CRL-2647) was purchased from ATCC (Manassas, VA, USA). Conditioned medium is prepared from each cell line grown and maintained in DMEM-F12 supplemented with 10% Corning FBS.
2. A second complete medium with growth factors (L-WRN medium), prepared at BCM by the DDC core, consisted of the same components as those of BCMp medium, with the exception that *N*-acetylcysteine was reduced by 50%, and Noggin, R-spondin and Wnt-3A were expressed from a single cell line, L-WRN ATCC CRL-3276 grown in DMEM-F12 supplemented with 20% Hyclone FBS (Manassas, VA, USA) following published information (36).
3. BCM differentiation medium (BCMd) consisted of the same components as those of BCMp medium without the addition of Wnt-3A, SB202190, and nicotinamide as well as 50% reduction in the concentrations of Noggin and R-spondin conditioned media. After 1 day of cell growth as a monolayer, the proliferation medium (BCMp or L-WRN) was changed to BCM differentiation medium. The cell monolayers were differentiated for 5 days as described above.
4. Commercial Intesticult (INT) human organoid growth medium (Stem Cell Technologies), composed of components A and B. The cell pellets, resulting from HIE cell dispersion, were suspended in proliferation INT medium (INTp), prepared by mixing equal volumes of components A and B, and supplemented with 10 μM ROCK inhibitor Y-27632.
5. After 1 day of cell growth as a monolayer, the INTp medium was changed with differentiation INT medium (INTd), consisting of an equal volume of component A and CMGF[-] medium. The cell monolayers were differentiated for 5 days as previously described.

### Human norovirus (HuNoV) infection of HIE monolayers

5-day differentiated HIE cell monolayers were washed once with CMGF[-] medium and inoculated with 5 μL HuNoV, diluted in 100 μL CMGF[-] medium supplemented with 500 μM GCDCA, for 1-2 hour at 37°C. The inoculum was removed and monolayers were washed twice with CMGF[-] medium to remove unbound virus. Differentiation medium (100 μL containing 500 μM GCDCA) was then added and the cultures were incubated at 37°C for 24 hours.

### RNA extraction

Total RNA was extracted from each infected well using the KingFisher Flex Purification System and MagMAX-96 Viral RNA Isolation Kit. RNA extracted at 1 hpi, was used as a baseline to determine the amount of input virus that remained associated with cells after washing the infected cultures to remove the unbound virus. Replication of virus was determined by RNA levels quantified from samples extracted at 24 hpi.

### Reverse Transcriptase Quantitative Polymerase Chain Reaction (RT-qPCR)

The primer pair and probe COG2R/QNIF2d/QNIFS (38) were used for GII genotypes and the primer pair and probe NIFG1F/V1LCR/NIFG1P (39) were used for GI.1 using qScript XLT One-Step RT-qPCR ToughMix reagent with ROX reference dye (Quanta Biosciences). Reactions were performed on an Applied Biosystems StepOnePlus thermocycler using the following cycling conditions: 50°C (15 min), 95°C (5 min), followed by 40 cycles of 95°C (15 sec) and 60°C (35 sec). A standard curve based on a recombinant HuNoV GII.4 [Houston virus (HOV)] or GI.1 (Norwalk virus) RNA transcript was used to quantitate viral genome equivalents (GEs) in RNA samples (40, 41). A 0.5 log_10_ increase in GEs after 24 hpi relative to the amount of genomic RNA detected at 1 hpi (after removal of the virus inoculum and two washes of the monolayers to remove unbound virus) was set as a threshold to indicate successful viral replication.

### Bile acid analysis

Media samples and sera were analyzed for bile acids using LC-MS/MS (Q-Exactive Orbitrap; Thermo Scientific) as described previously (42).

### Statistical analysis

Each experiment was performed twice, with three technical replicates of each culture condition and time point. Data from combined experiments are presented. All statistical analyses were performed on GraphPad Prism version 8.2.0 for Windows (GraphPad Software, La Jolla California USA). Samples with RNA levels below the limit of detection of the RT-qPCR assay were assigned a value that was one-half the limit of detection of the assay. Comparison between groups was performed using the Students t-test, with statistical significance determined using the Holm-Sidak method. P-values < 0.05 were considered statistically significant.

## Author contributions

K.E., R.L.A. and M.K.E. conceived and designed the study. K.E., N.W.C, V.R.T., B.V.A performed experiments; X-L.Z., F.H.N., Y.X. provided reagents/analytical tools. K.E., M.K.E., R.L.A., S.R., V.R.T. and S.E.C. analyzed data and wrote the paper, with all authors providing comments.

## Acknowledgments

This work was supported in part by Public Health Service grants AI-057788 (to M.K.E.), P30 DK 56338 and contract HHSN2722017000381 from the National Institutes of Health, by NIH P30 shared resource grant CA125123, NIEHS grants 1P30ES030285 and 1P42ES0327725, and by the John S. Dunn Research Foundation.

## Notes

### Competing Interest Statement

The authors have declared no competing interest.

